# TRACE: a framework for traceable biomedical resource construction demonstrated through single-cell glaucoma data

**DOI:** 10.64898/2026.11.04.764154

**Authors:** Xiaoqi Liang, Rojashree Jayakumar, Dario Strbenac, Marni Torkel, Beilei Bian, Eugenie Lee, Jasmine Gu, Vo Thu Nga Phan, Jonathan Crowston, Jean Yee Hwa Yang, Heejung Shim

**Affiliations:** Sydney Precision Data Science Centre, The University of Sydney, Sydney, NSW, Australia; School of Mathematics and Statistics, The University of Sydney, Sydney, NSW, Australia; Charles Perkins Centre, The University of Sydney, Sydney, NSW, Australia; Snow Vision Accelerator, The University of Sydney, NSW, Australia; School of Computer Science, The University of Sydney, NSW, Australia; Faculty of Medicine and Health, Save Sight Institute, The University of Sydney, Sydney, NSW, Australia; ARC Centre of Excellence for the Mathematical Analysis of Cellular Systems, The University of Sydney, NSW, Australia; School of Mathematics and Statistics, The University of Melbourne, MEL, Australia; Melbourne Integrative Genomics, The University of Melbourne, NSW, Australia

**Keywords:** Database, Glaucoma, Single-cell transcriptomics, Data curation

## Abstract

The growing availability of public omics datasets presents opportunities for biomedical research, yet constructing research question-driven resources from distributed studies remains challenging due to heterogeneous data, incomplete metadata and limited documentation of curation decisions. Here, we present TRACE (Traceable Resource Assembly and Curation Ecosystem), a general framework that links systematic knowledge discovery, data curation and resource delivery while preserving provenance throughout resource construction. We demonstrate TRACE through TRACE-scGlaucoma, a curated single-cell transcriptomics resource for glaucoma and optic nerve biology. Systematic screening identified 57 eligible studies, of which 42 provided accessible datasets comprising approximately 3.3 million cells. The resource provides harmonised metadata, standardised cell-type annotations and reusable expression objects while retaining original study information and curation provenance. We demonstrate its analytical utility through cellular enrichment of glaucoma GWAS-associated genes, comparing candidate drug-repurposing signals across distinct biological contexts and assessment of cell-type annotation reliability. TRACE provides a systematic and traceable approach to transforming fragmented public datasets into reusable biomedical resources, supporting reproducible analyses and future extensions across disease areas and data modalities.

## Introduction

Large-scale biological repositories have become essential resources for understanding cellular heterogeneity across tissues, diseases, and experimental conditions. Recent advances in omics technologies have led to the rapid expansion of publicly available repositories and atlas-scale datasets generated by international consortiums and independent studies^1 2^. These datasets are increasingly accessible through public repositories and data platforms, such as the Human Cell Atlas, Gene Expression Omnibus, CellxGene ^3 4 5^. They provide broad coverage across biological systems, experimental technologies, and disease contexts, enabling downstream applications in comparative analysis, biomarker discovery, and computational method development. However, translating this growing body of distributed public resources into a coherent, disease-focused resource still requires substantial dataset discovery, selection and curation ^6^.

Disease-focused data resources are commonly assembled through project-specific workflows involving keyword searches, manual publication screening and researcher-guided dataset selection ^78^. Relevant datasets are distributed across repositories, publications and supplementary materials, requiring manual extraction and organisation of associated metadata ^9^. Although the resulting collections may be valuable, their construction often involves considerable manual effort and researcher judgement in study selection, metadata interpretation and data organisation, with these decisions frequently incompletely documented. This limits the ability to audit how a resource was constructed, reproduce or update its contents consistently, and determine why potentially relevant datasets were excluded.

Existing repositories provide essential infrastructure for data deposition and access, but do not necessarily capture the full process through which a research question is translated into a curated, disease-focused data resource. Data provenance, which records the origins and transformation history of data, provides a foundation for tracking how datasets are generated and processed ^10^. However, constructing disease-focused resources requires traceability beyond individual datasets, encompassing decisions made during study identification, screening, selection and curation ^11^. An integrated framework connecting systematic study identification and screening with dataset acquisition, provenance tracking, metadata harmonisation, data processing and structured release would enable end-to-end traceability, improving the transparency, reproducibility and maintainability of biomedical data resources ^12 13^.

To address these challenges, we present TRACE, a Traceable Resource Assembly and Curation Ecosystem for constructing curated, disease-focused biomedical data resources ^14^. TRACE links systematic study identification and screening with dataset acquisition, metadata harmonisation, data processing and structured release, enabling each analysis-ready dataset to be traced back to its source and curation history. We implement TRACE through TRACE-scGlaucoma, a disease-focused resource for glaucoma and optic nerve biology. Its initial single-cell release comprises 57 eligible studies, 42 accessible datasets and around 3.3 million cells, together with harmonised metadata, reusable expression objects and documented processing workflows. The TRACE framework is designed to support subsequent incorporation of bulk and other biomedical data modalities.

### The TRACE-framework

TRACE (Traceable Resource Assembly and Curation Ecosystem) is a general framework for constructing disease-focused biomedical data resources from distributed public datasets. Rather than treating resource development as a collection of project-specific tasks, TRACE formalises the complete lifecycle of resource construction as a reproducible and traceable workflow. The framework comprises three interconnected components (Figure 1): knowledge discovery, which establishes the evidence base through systematic identification and assessment of relevant studies; knowledge curation, which transforms heterogeneous public datasets into reusable resources through standardised data processing and metadata harmonisation while preserving provenance; and knowledge delivery, which disseminates curated resources through structured, version-controlled releases that support exploration, reuse and future updates. Together, these components provide a general framework for translating distributed public datasets into traceable, reproducible and extensible biomedical resources. TRACE is independent of any specific disease area, biological modality or software implementation. In the following section, we demonstrate its application through TRACE-scGlaucoma, a disease-focused single-cell transcriptomics resource for glaucoma and optic nerve biology. Glaucoma is the leading cause of irreversible blindness worldwide. This optic neuropathy results from the injury and subsequent death of retinal ganglion cells that transmit the visual message from the retina to the brain.

**Figure 1.**
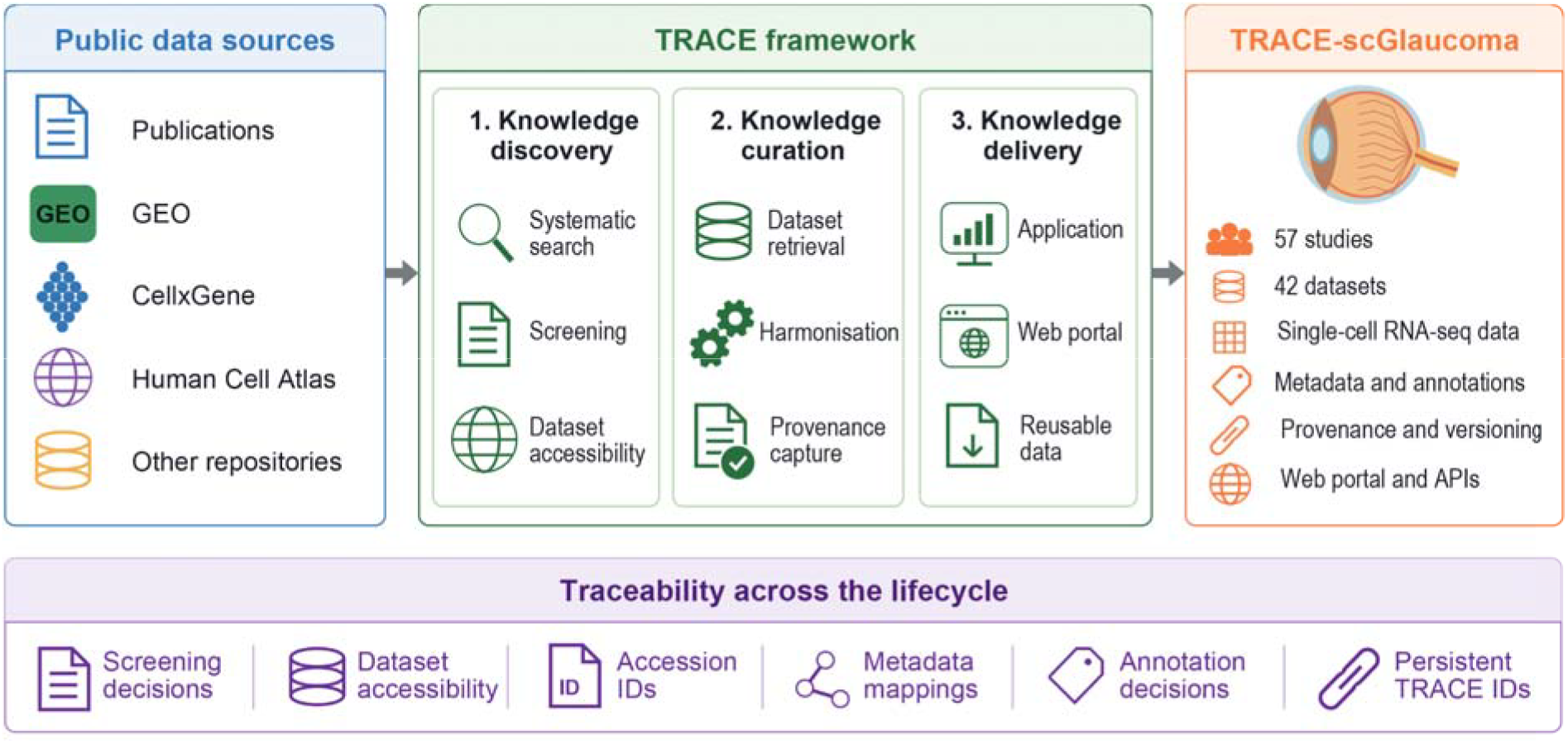
Overview of the TRACE framework. TRACE links knowledge discovery, knowledge curation and knowledge delivery within a unified workflow, maintaining traceability throughout the construction of disease-focused biomedical data resources from distributed public datasets.

### TRACE implementation through TRACE-scGlaucoma

#### Knowledge discovery

A systematic literature search was performed in Dec 2025 using the Ovid platform to identify single-cell transcriptomics studies related to glaucoma and optic nerve biology. The search strategy incorporated terms across four concepts: (i) single-cell transcriptomics; (ii) glaucoma diseases of interest; (iii) retinal cell types of interest (retinal ganglion cells and glia); and (iv) ageing-associated biological processes (ageing, mitochondrial and metabolic processes). The exact search parameters are provided in Table S1. A total of 562 records were identified, of which 114 duplicates were removed (113 by Covidence and one manually), leaving 448 records for title and abstract screening.

An overview of the study selection and dataset accessibility assessment workflow is provided in Figure S1. Title and abstract screening was conducted independently by two reviewers against prespecified eligibility criteria, with disagreements resolved through discussion and consensus. After title and abstract screening, 347 studies were excluded, leaving 101 studies for full-text assessment. Two reviewers independently reviewed full texts and recorded reasons for exclusion. A further 47 studies were excluded based on scientific eligibility criteria, leaving 54 eligible studies. Three additional glaucoma-related single-cell studies were identified through a manual Google search using single-cell keywords and included, resulting in a final cohort of 57 eligible studies (Table S2).

For each of the 57 eligible studies, we examined associated datasets, supplementary materials and accession identifiers to identify publicly available datasets suitable for reanalysis. Of these, 42 studies (73.7%) had accessible datasets suitable for reanalysis. The remaining 15 studies (26.3%) lacked reusable datasets: data were unavailable from listed repositories for six studies, data from corresponding authors were still awaited for eight, and potential data release after journal publication was anticipated for one. These studies were retained in the study catalogue, together with reasons for data unavailability, providing a traceable record of the knowledge discovery process.

#### Knowledge curation

##### Dataset acquisition and processing

Datasets associated with eligible studies were retrieved from their original public repositories using the accession identifiers reported in the corresponding publications and supplementary materials. The available datasets included raw sequencing files, gene-by-cell count matrices and, in some cases, author-processed expression objects. Datasets were downloaded from source repositories Gene Expression Omnibus, Zenodo, Mendeley, BioProject, Human Cell Atlas, ArrayExpress, and one study even hosts its Matrix Exchange files on GitHub, together with study- and sample-level metadata including accession identifiers, dataset title, tissue source, donor characteristics and experimental information when available.

Raw count matrices from the retrieved datasets were imported from 10x Genomics HDF5 or Matrix Market formats using DropletUtils version 1.32.0 or Matrix version 1.7-6. The imported data were stored as SingleCellExperiment objects to support standardised downstream analyses as described below ^15^. We removed likely empty droplets (<1000 UMI barcodes) when filtered data were unavailable. Between-cell normalisation was performed by the analyze.se function of scrapper version 1.6.3. Original study metadata and published cell-type annotations were retained to preserve consistency with the source datasets, where possible.

##### Metadata harmonisation and cell annotation

Metadata were extracted from repositories, publications and supplementary files and organised at the study, dataset, donor, sample and cell levels. Extracted metadata were manually verified by data curators. Variables included species, tissue, disease status, sequencing modality, donor or sample identifier, age, sex, timepoint, treatment, publication DOI and repository accession. Original field names and values were mapped to a standardised TRACE-scGlaucoma metadata schema.

Author-generated cell-type annotations were recovered where available through correspondence with study authors or matching cell barcodes to publicly deposited annotations. For datasets without recoverable author-generated annotations, cell identities were recorded as ‘Unshared’. Annotation availability, sources and modifications were documented for each dataset (Supplementary Methods: Recovery and assessment of cell-type annotations).

#### Knowledge delivery

##### Resource organisation

TRACE-scGlaucoma organises curated datasets into a structured resource that supports both reproducible analysis and efficient reuse. Each eligible study is represented through linked study-, dataset-, sample- and cell-level records, enabling biological observations to be traced across multiple levels of organisation (Figure S2). The processed data provide analysis-ready expression objects together with study-specific metadata (i.e., SingleCellExperimental objects) and standardised cell annotations.

Each study, dataset, sample and cell is ultimately assigned a persistent TRACE-scGlaucoma identifier that links metadata, expression objects, cell-type annotations and associated provenance records. This organisation enables users to recover the original study annotations while simultaneously performing comparative analyses using harmonised metadata and cell-type annotations.

##### Resource dissemination / Repository deposition and persistent identifiers

TRACE-scGlaucoma is publicly available through https://snow-vision-accelerator.github.io/TRACE-scGlaucoma/. The initial release comprises 57 eligible glaucoma-related studies, including 42 publicly accessible single-cell transcriptomics datasets, together with harmonised metadata, curated SingleCellExperiment objects, cell annotation tables and documented processing workflows. Source publications, repository accession numbers and provenance tables are retained throughout the resource to support traceability, reproducible reuse and future updates. Detailed descriptions of the database model, metadata schema, provenance table records and file organisation are provided in Supplementary Methods.

##### Interactive web interface

TRACE-scGlaucoma is accessed through an interactive web application that supports exploration of glaucoma-related single-cell transcriptomics datasets at the study, dataset and cellular levels (Figure S3). Users can browse datasets using biological and experimental metadata, including species, tissue, disease status and cell-type, and identify relevant datasets through searchable summaries and linked publication records. Individual datasets can be explored using interactive low-dimensional visualisations, where cells are coloured by harmonised or study-specific annotations, sample metadata and gene expression. Curated expression objects and harmonised metadata are available for download, while links to the original publications, repository accessions and provenance tables preserve traceability to the source data. The application is implemented using a React frontend communicating with a FastAPI backend, with additional implementation details provided in the Supplementary Methods (Figure 2) .

**Figure 2.**
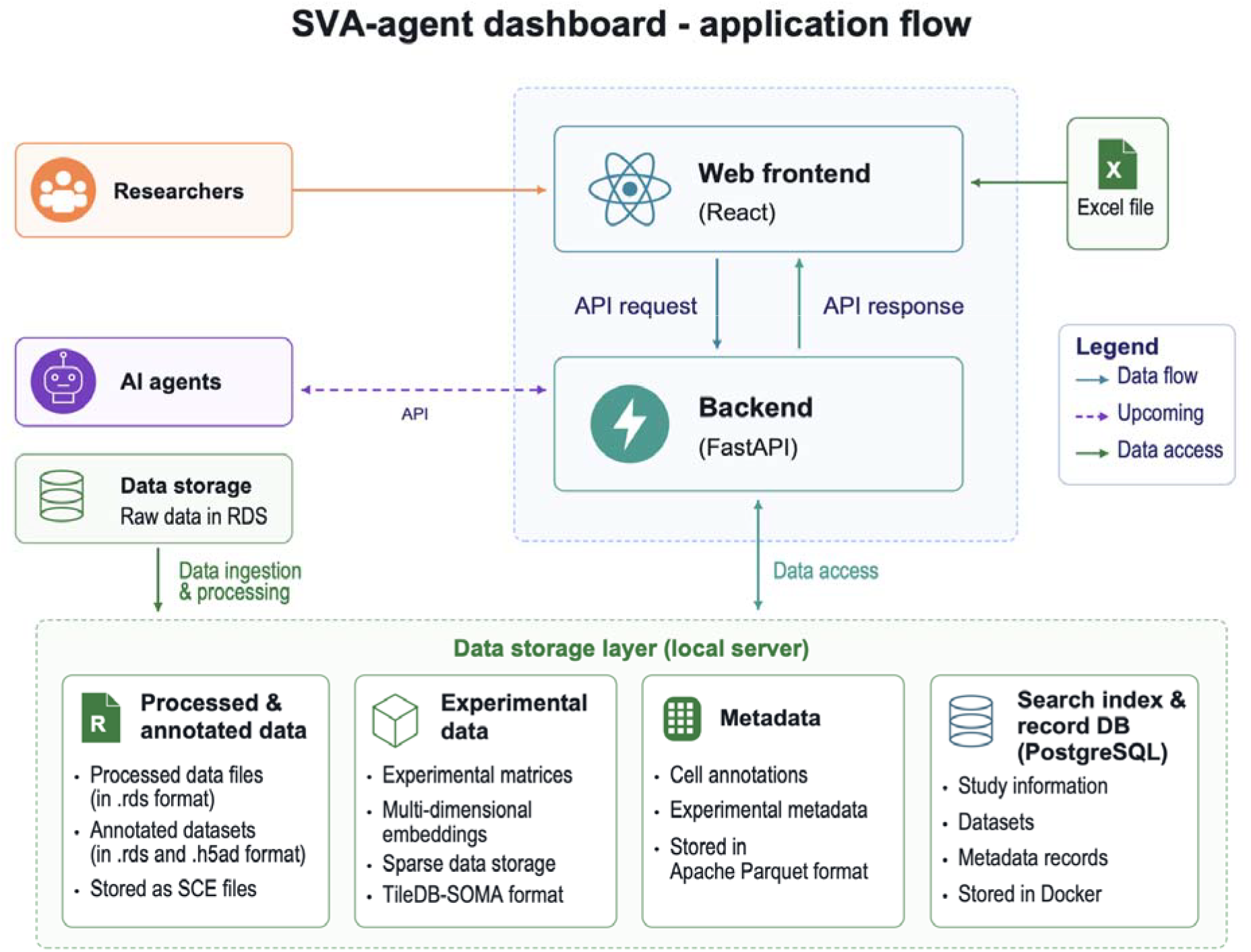
Architecture and data flow of the TRACE framework. Overview of TRACE architecture, showing the progression from the data storage layers through to structured data delivery and web implementation for glaucoma single-cell studies.

### Applications

TRACE-scGlaucoma includes datasets spanning different species, tissues, sequencing platforms and experimental designs, encompassing 88 annotated cell types across peripheral blood, retinal organoids, and anterior and posterior eye tissues. Given this biological and technical heterogeneity, analyses were conducted within individual datasets or compared across selected datasets, rather than combining all 42 datasets into a single integrated low-dimensional embedding. To illustrate the analytical utility of TRACE-scGlaucoma, we present three applications: assessing the reliability of cell-type annotations, examining the cellular enrichment of glaucoma-associated genes and comparing drug-repurposing predictions across distinct biological contexts.

#### Assessing cell-type annotation reliability

Cell-type annotation is essential for downstream single-cell analyses, yet its reliability can be affected by analytical choices and ambiguity in cell identity ^16^. To demonstrate how TRACE-scGlaucoma can support the assessment of annotation reliability, we evaluated the stability of cell-type assignments in dataset D002 under analytical perturbations (see Supplementary Methods). Per-cell uncertainty UMAP showed that most cells retained a stable label, with uncertain cells concentrated at the edges of specific clusters rather than spread evenly across the dataset (Figure 3A). However, uncertainty varied across cell populations (Figure 3B). Müller glia showed the highest mean uncertainty (7.5%), followed by smooth muscle and cone bipolar cells (6.4% each). In contrast, rods (0.7%), melanocytes (1.5%), retinal pigment epithelium (1.6%) and uveal fibroblasts (1.7%) exhibited lower uncertainty. Comparison of mean annotation uncertainty with cell-type proportion showed variability among less abundant cell populations, while the most abundant cell types exhibited lower uncertainty (Figure S4). These results illustrate that annotation stability varies across cell populations, highlighting the value of quantifying cell-level uncertainty to inform the interpretation and robustness assessment of downstream analyses.

**Figure 3.**
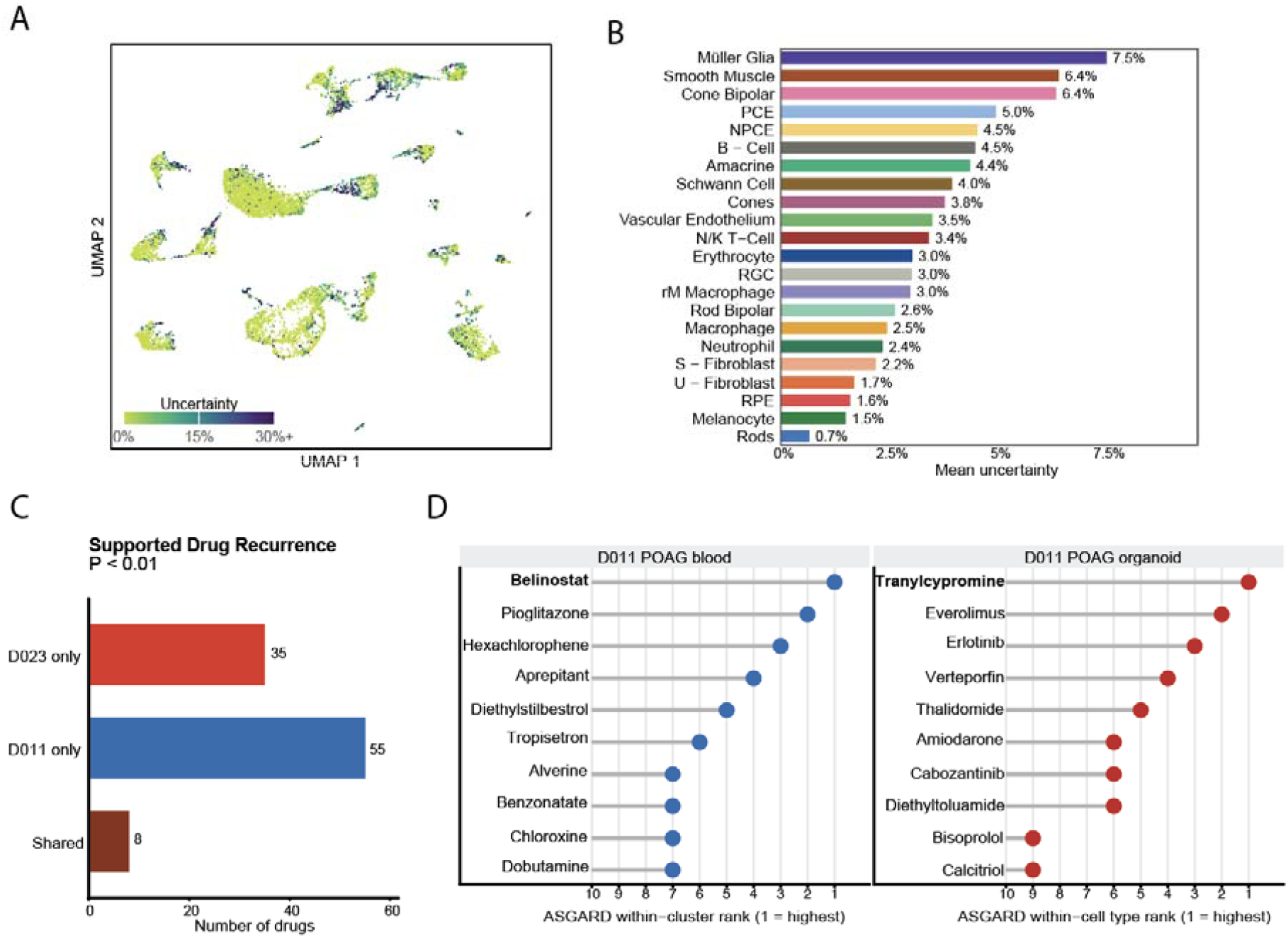
Drug-repurposing analysis and cell-type annotation uncertainty in TRACE-scGlaucoma. (A) Number of supported drug candidates shared between or specific to the two datasets, defined by a minimum ASGARD P value < 0.01 across cell types or clusters. (B) Top 10 ASGARD-ranked drug candidates in POAG blood (D011) and organoid (D023) datasets. (C) UMAP showing per-cell annotation uncertainty in D002. (D) Mean annotation uncertainty by cell type in D002.

#### Cross-dataset comparison of candidate drug-repurposing signals

To illustrate the utility of TRACE-scGlaucoma for downstream drug-repurposing analyses, we applied ASGARD, a computational framework that prioritises candidate drugs based on their potential to reverse disease-associated gene-expression changes ^17^. We selected two independent POAG datasets with case-control contrasts, D011 (blood) and D023 (iPSC-derived retinal organoids), representing distinct biological contexts. Using a minimum ASGARD P value < 0.01 across cell clusters or types within each dataset, we identified 8 candidate drugs supported in both datasets, alongside 35 and 55 candidates identified only in D011 and D023, respectively (Figure 3C). The highest-ranked candidates also differed between datasets, with belinostat prioritised in D011 and tranylcypromine in D023 (Figure 3D). Both candidates have plausible biological relevance to retinal neuroprotection, with previous studies implicating histone deacetylase (HDAC) inhibition and lysine-specific demethylase 1 (LSD1) inhibition in retinal ganglion cell protection ^18 19^. Together, these findings illustrate how TRACE-scGlaucoma supports the application of existing analytical methods to curated datasets, enabling comparisons of shared and distinct drug-repurposing predictions and the identification of biologically plausible candidates for further investigation.

#### Cellular enrichment of glaucoma GWAS-associated genes

To demonstrate how TRACE-scGlaucoma supports integrative analysis with external genomic resources, glaucoma GWAS signals were mapped onto two single-cell datasets representing complementary biological contexts, including healthy adult retina (D024) and patient-derived retinal organoids (D023) (see Methods). Figure 4 shows the disease relevance scores that were calculated for individual cells using scDRS ^20^ and summarised across annotated cell populations. In the healthy adult retina, significant group-level associations were identified in vascular, pigmented and macroglial populations (P = 0.001), with 80.8%, 60.2% and 15.2% of cells showing significant cell-level disease relevance scores (FDR < 0.1), respectively (Figure 4A-B, Table S3). In the primary open-angle glaucoma (POAG) iPSC-derived retinal organoid, glaucoma-associated signatures were preferentially detected in specific retinal progenitor populations, particularly RPC4 (P = 0.002), which also exhibited the highest proportion of significant cells (2.5%, FDR < 0.1; Figure 4C-D, Table S4). These analyses illustrate how TRACE-scGlaucoma can support cell-level mapping of glaucoma-associated gene-expression signals in two datasets representing distinct developmental and disease contexts.

**Figure 4.**
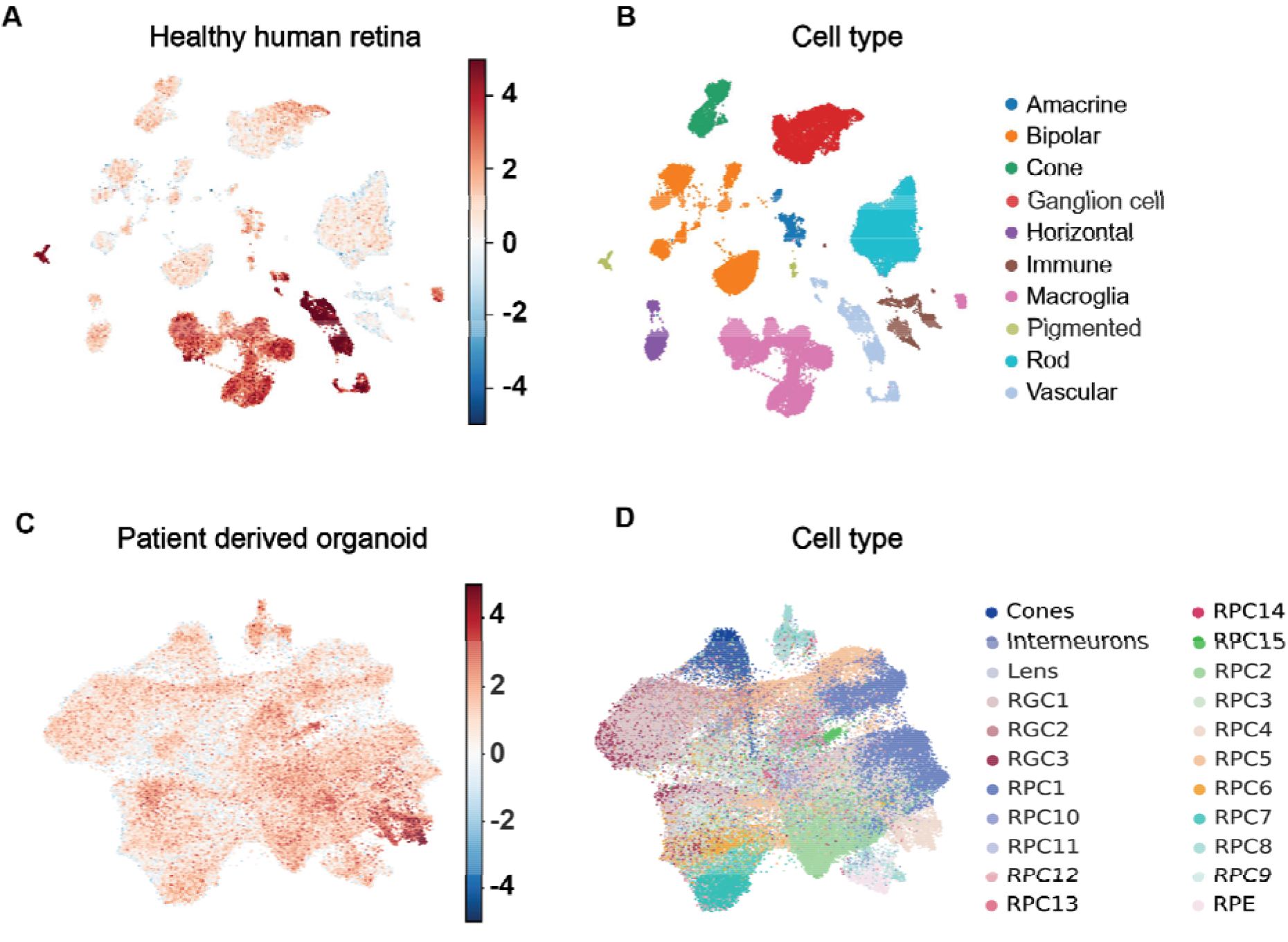
Cellular enrichment of glaucoma GWAS-associated genes across healthy adult retina and patient-derived retinal organoids. (A) UMAP of a healthy adult human retinal single-cell RNA-seq dataset coloured by scDRS glaucoma disease-relevance scores, with higher scores indicating greater expression of genes associated with glaucoma genetic risk. (B) Corresponding UMAP showing retinal cell-type annotations. (C) UMAP of a primary open-angle glaucoma (POAG) iPSC-derived retinal organoid dataset coloured by scDRS glaucoma disease-relevance scores. (D) Corresponding UMAP showing retinal progenitor cell (RPC) and differentiated retinal cell annotations.

## Discussion

In this study, we introduced TRACE, a general framework for constructing traceable and reproducible disease-focused biomedical resources, and demonstrated its implementation through TRACE-scGlaucoma. By linking systematic knowledge discovery, data curation and resource delivery, TRACE enables the documentation of decisions throughout resource construction. A key contribution of TRACE is extending traceability across resource construction, from study identification, screening and selection decisions to data acquisition, including the source and accessibility of datasets, processing strategies and the provenance of curated data. The TRACE-scGlaucoma implementation demonstrates this approach through the systematic identification of 57 eligible studies, of which 42 provided accessible datasets suitable for reanalysis.

Beyond resource construction, TRACE-scGlaucoma demonstrates how systematically curated datasets can support biomedical analyses across different biological contexts. Applications included localisation of glaucoma-associated genetic risk across retinal cell populations, comparison of candidate drug-repurposing signals across independent datasets, and assessment of cell-type annotation uncertainty. Together, these applications illustrate how a traceable, curated resource can support cross-dataset investigations, comparison of disease-associated signals and assessment of annotation reliability.

Biomedical data curation is not merely a technical preprocessing step but can influence downstream scientific inference. In TRACE-scGlaucoma, only 42 of 57 eligible studies provided accessible datasets suitable for reanalysis, illustrating how data accessibility shapes the evidence available for investigation. Incomplete metadata and cell-type annotations can further constrain data reuse and biological interpretation ^21 22 23^, while variation in annotation reliability may influence downstream cell-type-specific analyses. By documenting dataset accessibility, annotation availability and associated curation decisions, TRACE makes these factors more transparent, allowing researchers to assess how resource construction may influence subsequent analyses.

The construction of TRACE-scGlaucoma highlighted opportunities for improving biomedical data curation. Recovering cell-type annotations required searching external repositories, contacting authors and matching cell identifiers, while reproducing unavailable annotations involved additional analytical decisions (see Methods). These challenges illustrate the need for expert judgement in interpreting incomplete biological information, which becomes increasingly difficult to scale as public datasets grow. While AI-assisted workflows could improve literature surveillance, metadata extraction and annotation recovery, potentially reducing manual effort^24 25 26^, their accuracy and consistency remain variable across tasks and models ^27 28^. Validation, expert oversight and documentation of automated decisions and their provenance are therefore essential for supporting reliability and reproducibility.

Despite these contributions, TRACE-scGlaucoma remains subject to limitations associated with the scope and heterogeneity of publicly available datasets. Included studies vary in species, tissue source, experimental design, sequencing technology and metadata completeness, introducing biological and technical heterogeneity that cannot be resolved through metadata and annotation standardisation ^29^. Although harmonised metadata and organised cell-type annotations support comparative and exploratory analyses across datasets, these differences must be considered when interpreting results. Preserving original expression matrices, metadata and study-specific annotations is therefore important for supporting reproducibility and study-specific interpretation. In addition, TRACE-scGlaucoma does not capture unpublished or restricted-access datasets that may provide additional biological contexts relevant to glaucoma research.

While TRACE-scGlaucoma focuses on single-cell transcriptomic datasets, the TRACE framework could be extended to support biomedical resources incorporating additional data modalities, including bulk transcriptomics, spatial transcriptomics, spatial proteomics and imaging ^30^. Such extensions would require modality-specific curation and processing strategies while preserving common principles of systematic knowledge discovery, provenance tracking and resource delivery.

## Conclusion

TRACE provides a general framework for constructing traceable and reproducible biomedical data resources. Through TRACE-scGlaucoma, we demonstrated how linking systematic knowledge discovery, data curation and resource delivery supports the construction and reuse of curated single-cell datasets. By documenting decisions from study identification and selection through to data acquisition, processing and provenance, TRACE establishes a foundation for transparent and reproducible biomedical resource construction.

## BACKMATTER

### Ethics approval and consent to participate

Not applicable

### Consent for publication

Not applicable

### Data Availability

Our curated datasets for the publications can be downloaded through https://snow-vision-accelerator.github.io/TRACE-scGlaucoma/.

### Code Availability

The analysis scripts and source data are available on GitHub at https://github.com/Snow-Vision-Accelerator/TRACE-scGlaucoma.

## Acknowledgments

The authors thank all their colleagues, particularly at The University of Sydney, Sydney Precision Data Science and Charles Perkins Centre for their support and intellectual engagement. We also thank Paul Szabo and the University of Sydney ICT team for their technical assistance and support.

## Author Contribution

J.Y.H.Y. and H.S. conceived and led the study. B.B., E.L, J.G., J.Y.H.Y., R.J. V.T.N.P and X.L. conducted the literature search and study screening. B.B., E.L, R.J. and X.L. acquired and processed datasets. D.S. curated the datasets, metadata and cell-type annotations. M.T organised the resource and developed the web platform. B.B., D.S., E.L. and X.L. ran the case studies and performed applications. The development of the designed template was done jointly by all authors and all authors wrote, reviewed and approved the manuscript.

## Supplementary Methods

### Knowledge curation

#### Raw, processed and curated data layers

Data is organised into two linked layers. The source-data layer records the original repository accession, available file types and source location for each dataset. The resource provides persistent links to the source repository. The curated-data layer contains, standardised study-, donor-, sample- and cell-level metadata, and cell-type annotations. Integrated representations are stored separately from the source count matrices and should not be interpreted as replacements for the original data.

#### Metadata tables

The resource contains linked metadata tables describing studies, datasets, donors, samples and cells. The study table records publication-level information, including TRACE-scGlaucoma study identifier, title, DOI, publication year, eligibility status and source repository. The dataset table records accession identifiers, species, tissue, disease context, sequencing modality, available data layer and download status. Donor- and sample-level tables contain available demographic, biological and experimental variables, including age, sex, disease status, tissue source and sample identifiers. Cell-level metadata include harmonised cell annotations, quality-control variables and links to the corresponding dataset and sample.

#### Recovery and assessment of cell-type annotations

##### Recovery of author-generated cell-type annotations

During dataset curation, we assessed the availability of author-generated cell-type annotations for each study. Although expression count matrices were accessible for many datasets, the corresponding cell-type labels were mostly unavailable. We recovered annotations for five studies (D003, D007, D021, D041, and D042) through correspondence with the corresponding authors. For a further seven studies (D012, D018, D019, D020, D033, D034 and D037), annotations were retrieved from the Broad Institute Single Cell Portal and matched to the expression matrices using cell barcodes. Twelve studies remained without accessible author-generated cell-type annotations.

##### Reproduction of published cell-type annotations

Reproduction is challenging when original analytical code or processing parameters are unavailable. For example, initial reanalysis of dataset D002 using default clustering parameters identified 22 clusters, matching the number reported in the original publication. However, the resulting cluster structure and marker-gene patterns did not consistently reproduce the published annotations. To avoid introducing potentially divergent cell-type assignments, we did not systematically assign new annotations to datasets for which author-generated labels could not be reliably recovered or reproduced.

### Knowledge delivery

#### File structure, relationships and provenance

Each study, dataset, donor, sample and cell is assigned a persistent TRACE-scGlaucoma identifier. These identifiers provide the keys linking the metadata tables to the corresponding expression objects. The relationship among these objects is summarised in Figure S2.

A provenance table records the source publication and repository accession for each released object, together with the original input files, metadata transformations, annotation source and TRACE-scGlaucoma release version. This enables users to trace each curated dataset from the final released object back to its original study and repository record. All datasets can be downloaded though https://snow-vision-accelerator.github.io/TRACE-scGlaucoma/.

#### Web interface implementation

TRACE-scGlaucoma is implemented as a web-based platform comprising a React frontend and a FastAPI backend. User interactions, including dataset selection, metadata filtering and visualisation requests, are transmitted to the backend, which retrieves the corresponding metadata and expression summaries from the underlying database before returning formatted data for display.

TRACE-scGlaucoma uses a PostgreSQL catalogue to organise dataset descriptions, biological classifications, publication links and references to associated data assets. Dataset-, cell- and feature-level metadata are stored in Apache Parquet, while processed expression matrices and their cell and feature annotations are stored in TileDB-SOMA. Dataset identifiers and asset locators connect catalogue records to these files, with cell identifiers linking cell-level metadata to precomputed PCA, UMAP and t-SNE coordinates. Interactive visualisations use precomputed UMAP and t-SNE embeddings, with WebGL rendering supporting responsive exploration of large datasets. Current query times are approximately one or two seconds for datasets containing around 10,000 cells

The web interface provides two principal components:

- Overview summarises database composition, including the numbers of studies, cells and cell types, species distributions and publication timelines. It also describes the harmonised data and provides options for downloading and exploring datasets.
- Explore enables interactive visualisation of individual datasets, with cells coloured by original or harmonised annotations, sample metadata or selected gene expression. Users can filter cells by cell type, age and sex where these annotations are available.

The complete software environment, database reconstruction workflow and associated scripts are maintained under version control. Database snapshots are archived to support reproducibility and disaster recovery. Future releases will include documented programmatic APIs and additional indexing strategies to further improve scalability for larger atlas-scale resources.

### Analytical applications

#### Reliability calculation in cell-type annotation using perturbation

Count matrices for the mouse ciliary body dataset were obtained from GEO (GSE178667) ^31^. The published cell type labels were recovered by reproducing the original Seurat analysis (v5.3.0) ^32^: cells with 200–6,000 detected genes, under 25% mitochondrial reads and under 25,000 UMIs were retained (10,024 cells) and clustered on 2,000 variable genes and 25 principal components at a resolution of 0.3, yielding the 22 published clusters. Reliability was estimated with four reference-based annotation tools: Seurat label transfer, SingleR (v2.10.0) ^33^, CellTypist (v1.7.1, with a fixed random seed) ^34^ and scClassify (v1.20.0) ^35^. Each tool was trained on one of the two experiments with its published labels and used to label the other, and vice versa. The query cells were then perturbed by retaining 90% or 80% of their UMIs through random downsampling, or by removing a random 10% or 20% of genes, with five replicates of each, and relabelled by all four tools. For each cell, reliability was defined as the percentage of its 80 perturbed predictions (4 tools × 4 perturbations × 5 replicates) that differed from the label assigned by the same tool to the unperturbed cell, with unassigned calls counted as a distinct outcome. Cell-type reliability was the mean of the per-cell values. Analyses were performed in R (v4.5.1) and Python (v3.11).

#### ASGARD-Based Drug Repurposing Analysis

ASGARD was applied to two independent primary open-angle glaucoma (POAG) case-control datasets curated in TRACE-scGlaucoma: D011, a POAG blood dataset, and D023, a POAG organoid dataset ^17^. For each dataset, POAG and control cells were compared within available cellular units. For D023, curated cell-type annotations were used. For D011, where cell-type annotations were unavailable, cells were grouped by unsupervised transcriptomic clusters. Within each cell type or cluster, differential expression between POAG and control cells was performed using limma ^36^. Genes were ranked by the resulting test statistic and used as cluster- or cell-type-specific disease signatures for ASGARD.

Drug-reference profiles were generated from LINCS L1000 datasets GSE70138 and GSE92742 ^37^. In the absence of an eye-specific LINCS reference, the central nervous system reference was used. ASGARD was run using negative connectivity to identify drugs predicted to reverse POAG-associated transcriptional signatures. ASGARD generated an independent ranked drug list for each cellular unit, with lower ranks indicating stronger predicted reversal. Repeated occurrences of a drug across clusters or cell types were collapsed, and its dataset-level rank was defined as its lowest observed ASGARD rank across those units. Supported candidate drugs were defined as drugs with a minimum ASGARD P value < 0.01 across clusters or cell types. Recurrence was assessed by comparing supported candidates between D011 and D023 and classifying drugs as shared or dataset-specific.

#### Cellular enrichment of glaucoma GWAS-associated genes

Genome-wide association study (GWAS) summary statistics for primary open-angle glaucoma (POAG) were obtained from individuals of European ancestry in the VA Million Veteran Program ^38^. Gene-level association statistics were generated using MAGMA with default settings and used as disease gene weights for downstream single-cell analyses ^39^. Two retinal single-cell RNA sequencing datasets from TRACE-scGlaucoma were selected to represent complementary biological contexts. The first comprised healthy adult human retina, including major retinal cell types, while the second comprised induced pluripotent stem cell (iPSC)-derived retinal organoids from glaucoma donors, representing developmental retinal cell states associated with disease ^40 41^. Both datasets were processed using the standard TRACE-scGlaucoma workflow, including quality control and normalisation.

Cell-level enrichment of glaucoma GWAS-associated gene expression was evaluated using scDRS, with MAGMA-derived gene-level association statistics supplied as disease gene weights ^20^. scDRS calculates a disease relevance score for each cell by comparing the expression of disease-associated genes with matched control gene sets while accounting for gene-specific technical variation.

Disease relevance scores were projected onto the UMAP. For each dataset, the proportion of disease-associated cells within each annotated cell type was calculated using the default scDRS significance threshold (FDR < 0.1).

