## Supplementary Methods and Figures for "TRACE: a framework for traceable biomedical resource construction demonstrated through single-cell glaucoma data"

### **Supplementary Tables**

**Supplementary Table S1: Complete search strategy for TRACE-scGlaucoma.** The literature search was conducted using the Ovid platform to identify single-cell and single-nucleus transcriptomic studies relevant to glaucoma and optic nerve biology. Search records were exported to Covidence for screening. Drug-related studies were excluded unless they contained relevant single-cell or single-nucleus transcriptomic data.

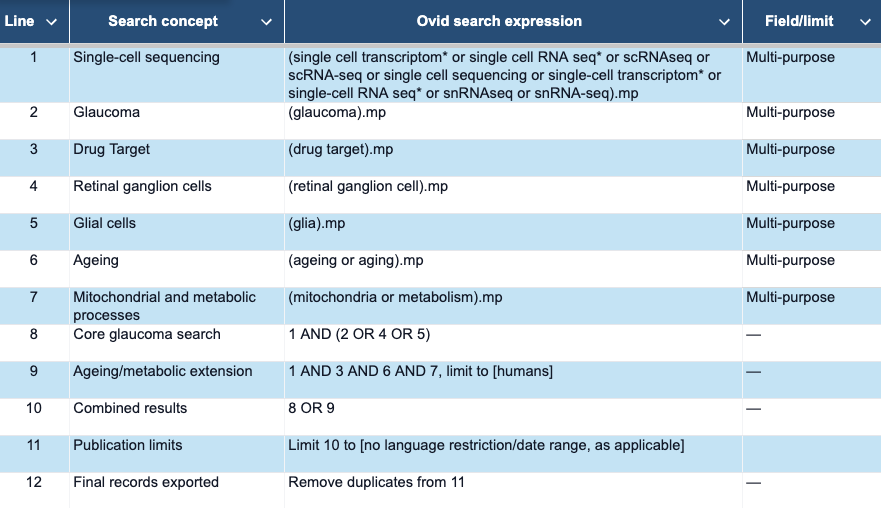

**Supplementary Table S2: Summary of datasets identified for TRACE-scGlaucoma**, including data sources, study details, sample and cell counts, tissues, diseases, assay platforms, and cell-type annotation sources.

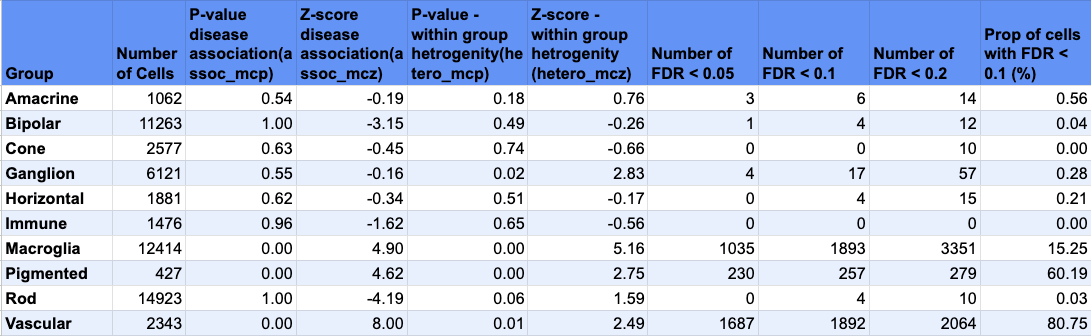

**Supplementary Table S3:** Cell-type-level association and heterogeneity analysis of glaucoma-associated genes in the healthy adult retina using scDRS.

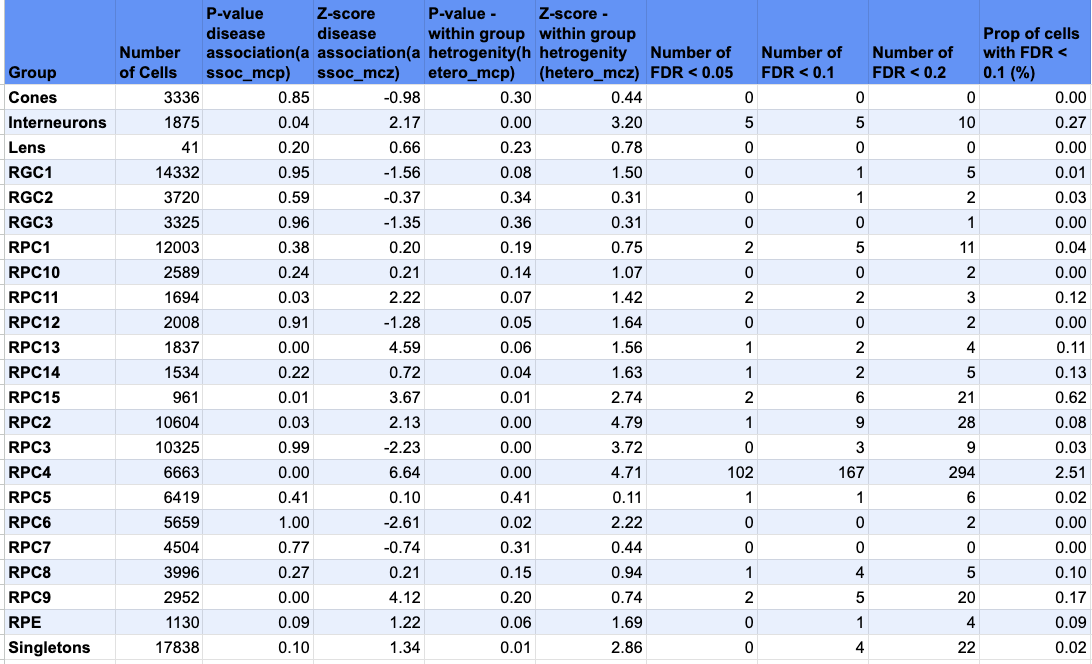

**Supplementary Table S4:** **Cell-type-level association and heterogeneity analysis of glaucoma-associated genes in patient-derived retinal organoids using scDRS**. RGC stands for Retinal Ganglion Cells; RPC stands for Retinal Progenitor Cells; RPE stands for Retinal Pigmented Epithelial

**Supplementary Figure**

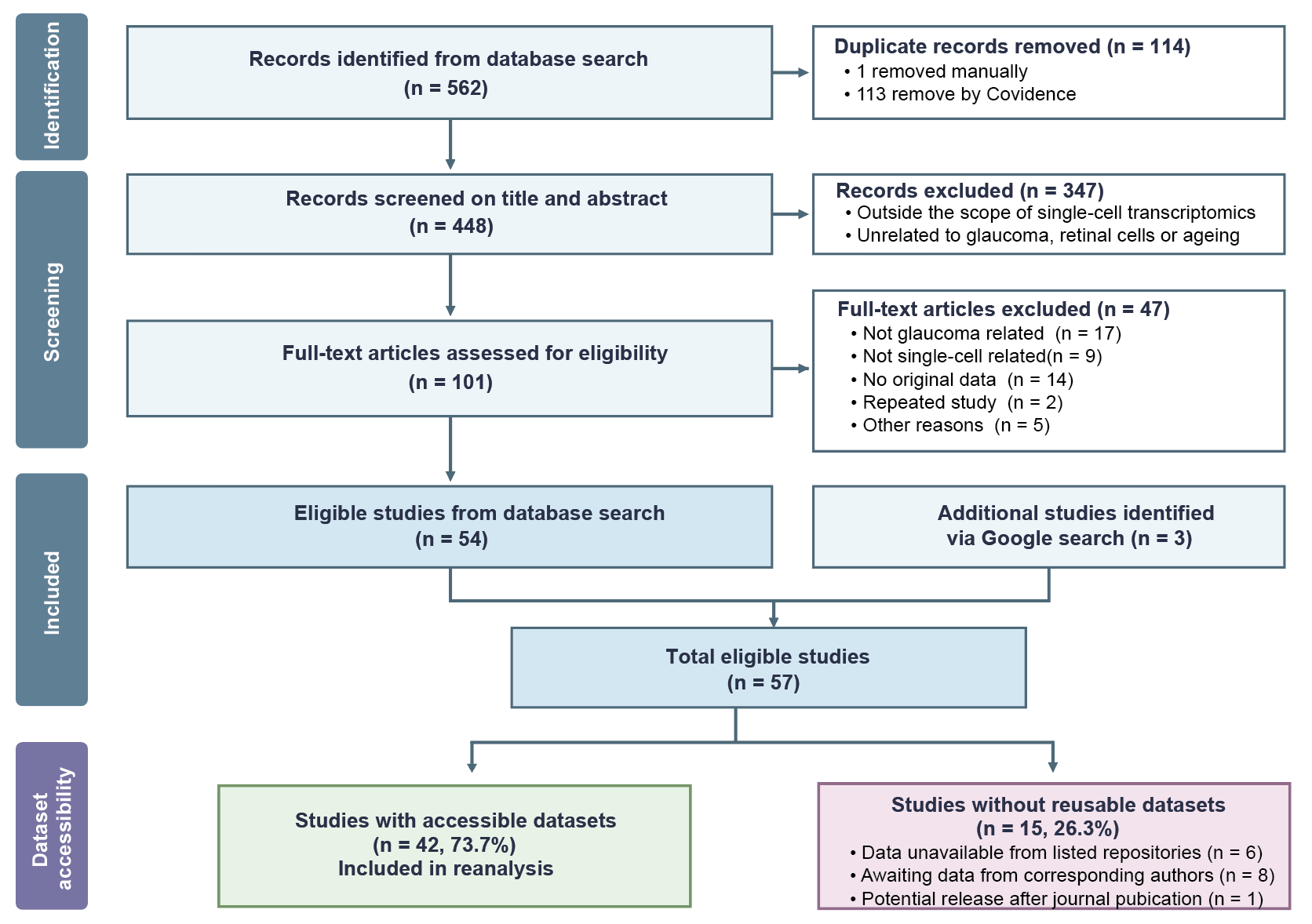

**Supplementary Figure S1:**
Systematic review workflow and dataset availability assessment for glaucoma single-cell studies.

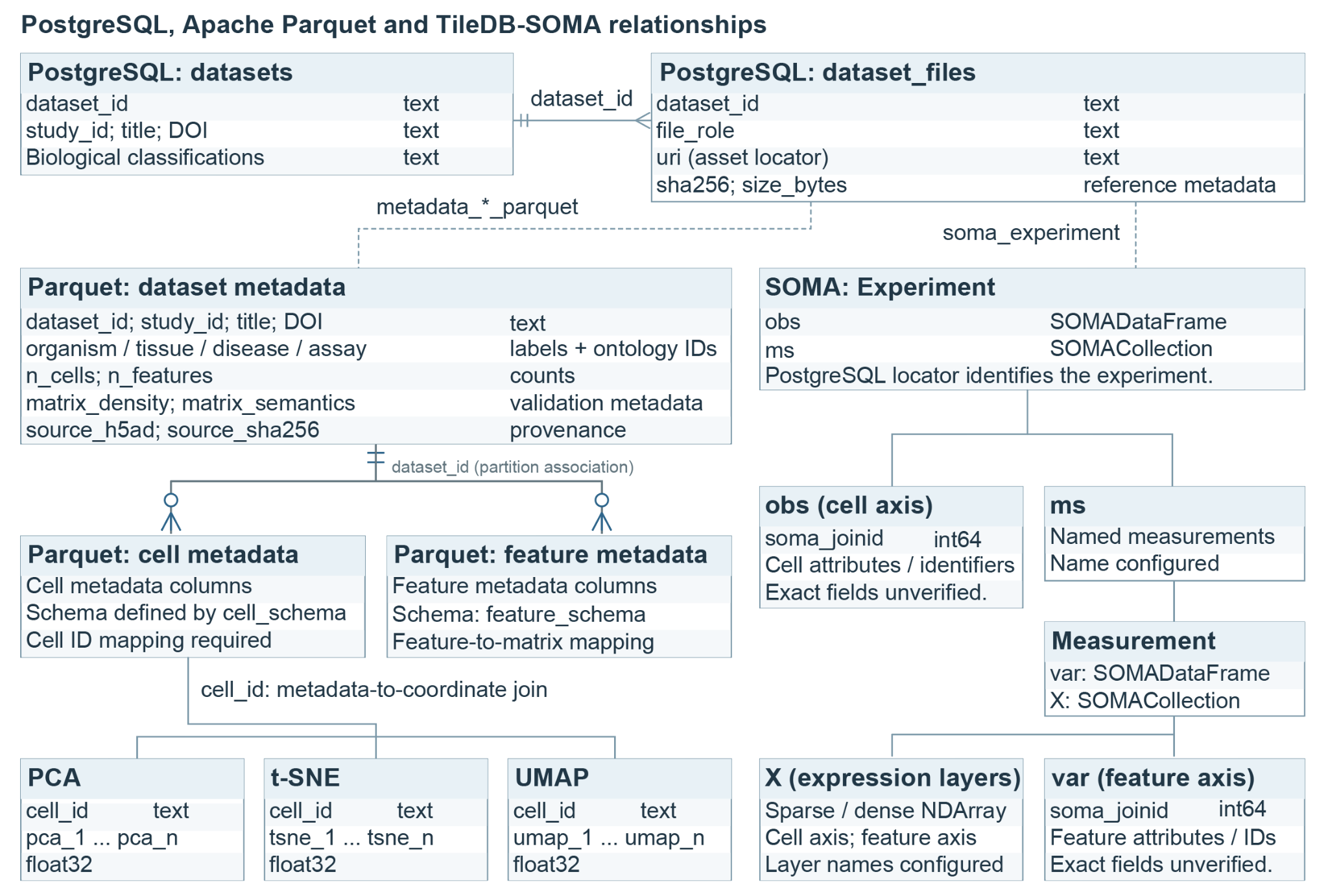

**Supplementary Figure S2: Data structure schema of TRACE-scGlaucoma.**
Relationships linking studies, metadata and data layers to preserve provenance and support reuse.

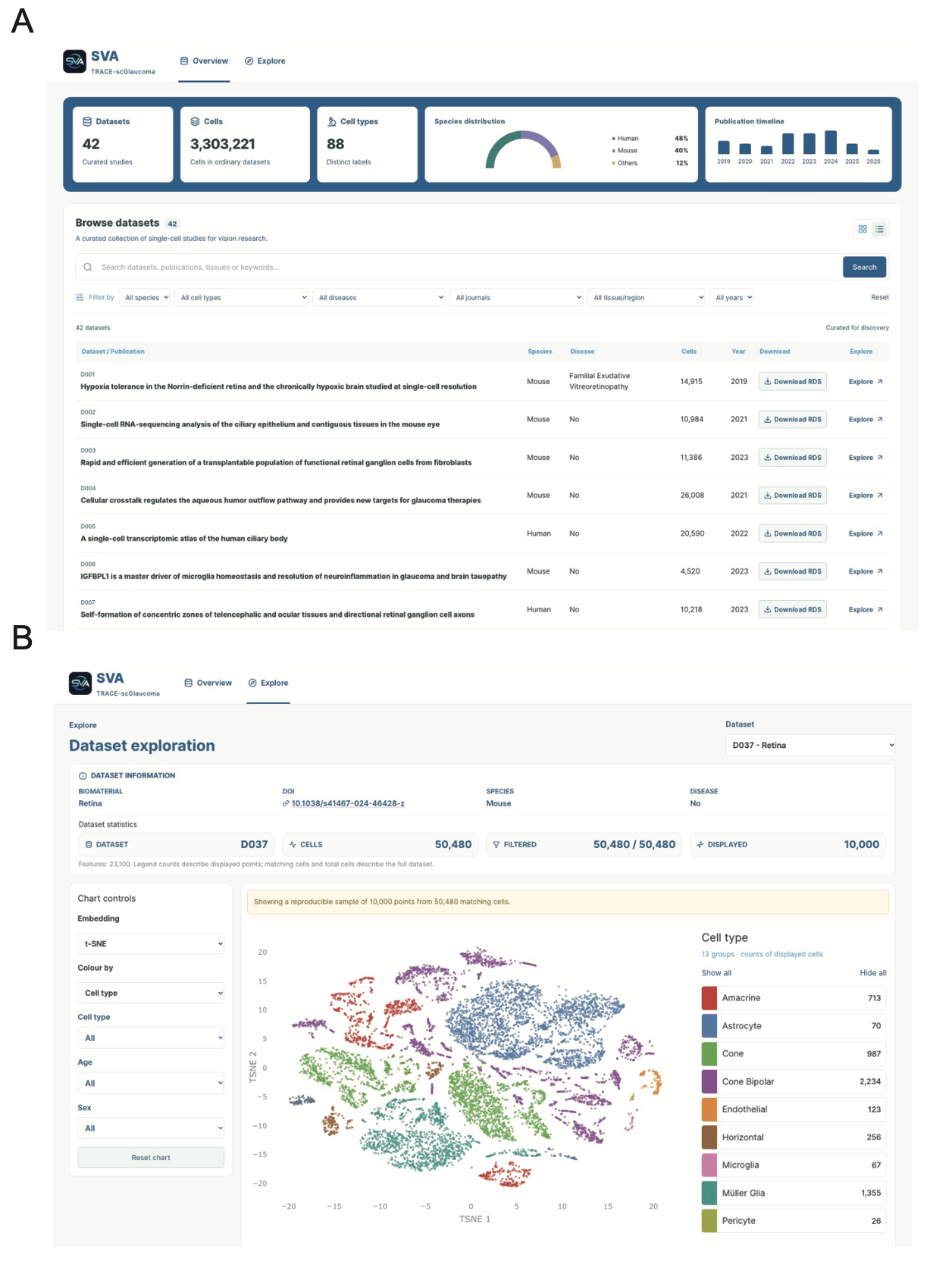

**Supplementary Figure S3: Overview of the TRACE-scGlaucoma web interface.** (A) Dataset overview and browsing interface. (B) Interactive exploration of example dataset D037, featuring metadata filtering and cell-level visualisation using t-SNE. The 88 annotated cell types reflect the diversity of tissues and biological systems represented, including peripheral blood, retinal organoids, and anterior and posterior eye tissues.

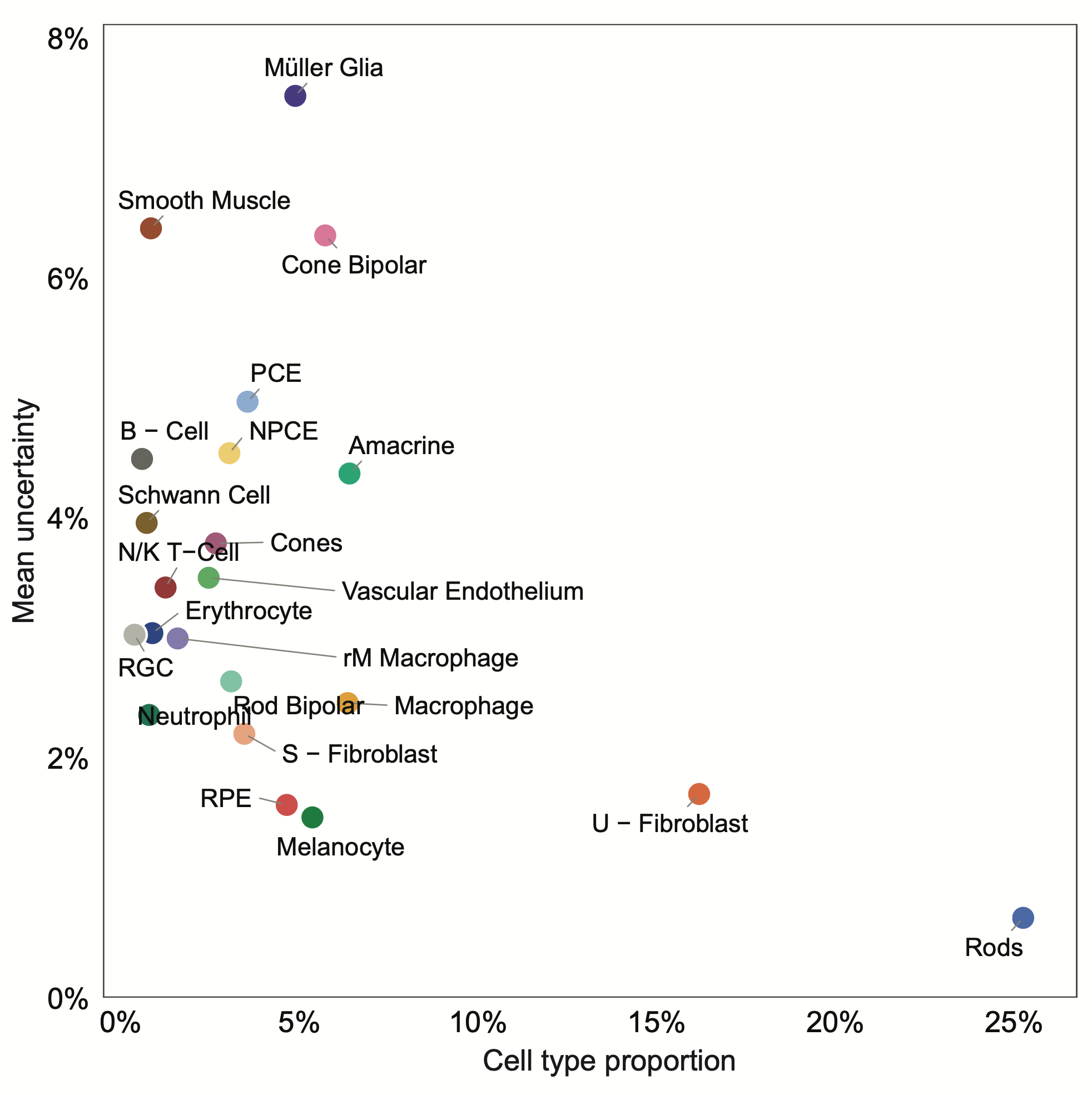

**Supplementary Figure S4: Cell type proportion against mean annotation uncertainty.**
Mean annotation uncertainty of each of the 22 published cell types plotted against the proportion of all 10,024 cells that belong to that type. Each point represents one cell type. Per-cell uncertainty was defined as the percentage of 80 perturbed predictions that differed from the original cell type label for the unperturbed cell (see Supplementary Methods).
